# A simple magnetic nanoparticles-based viral RNA extraction method for efficient detection of SARS-CoV-2

**DOI:** 10.1101/2020.02.22.961268

**Authors:** Zhen Zhao, Haodong Cui, Wenxing Song, Xiaoling Ru, Wenhua Zhou, Xuefeng Yu

**Affiliations:** Materials and Interfaces Center, Shenzhen Institutes of Advanced Technology, Chinese Academy of Sciences, Shenzhen 518055 (P. R. China); University of Chinese Academy of Sciences, Beijing 100049 (P. R. China)

## Abstract

The ongoing outbreak of the novel coronavirus disease 2019 (COVID-19) originating from Wuhan, China, draws worldwide concerns due to its long incubation period and strong infectivity. Although RT-PCR-based molecular diagnosis techniques are being widely applied for clinical diagnosis currently, timely and accurate diagnosis are still limited due to labour intensive and time-consuming operations of these techniques. To address the issue, herein we report the synthesis of poly (amino ester) with carboxyl groups (PC)-coated magnetic nanoparticles (pcMNPs), and the development of pcMNPs-based viral RNA extraction method for the sensitive detection of COVID-19 causing virus, the SARS-CoV-2. This method combines the lysis and binding steps into one step, and the pcMNPs-RNA complexes can be directly introduced into subsequent RT-PCR reactions. The simplified process can purify viral RNA from multiple samples within 20 min using a simple manual method or an automated high-throughput approach. By identifying two different regions (*ORFlab* and *N* gene) of viral RNA, a 10-copy sensitivity and a strong linear correlation between 10 and 10^5^ copies of SARS-CoV-2 pseudovirus particles are achieved. Benefitting from the simplicity and excellent performances, this new extraction method can dramatically reduce the turn-around time and operational requirements in current molecular diagnosis of COVID-19, in particular for the early clinical diagnosis.

## 2 Introduction

In December, 2019, an unknown pathogen-mediated pneumonia emerged in Wuhan, China.^1^ Confirmed cases had been soon diagnosed in other cities in China, as well as in other countries. On 11 February 2020, as reported by the World Health Organization (WHO) and the Chinese Center for Disease Control and Prevention (China CDC), this unknown pneumonia was confirmed to be caused by a novel coronavirus (SARS-CoV-2, previously named as 2019-nCoV). SARS-CoV-2 is etiologically related to the well-known severe acute respiratory syndrome coronavirus (SARS-CoV), belonging to the *Coronaviridae* family, a type of positive-sense, single-stranded RNA coronavirus with an outer envelope. Although the genome sequences of SARS-CoV-2 have been fully revealed and various RT-PCR-based detection kits have been developed, clinical diagnosis of COVID-19 is still highly challenging.

Due to its high sensitivity in exponentially amplifying RNA molecules, reverse transcription polymerase chain reaction (RT-PCR) is identified as a standard and routinely used technique for the analysis and quantification of various pathogenic RNA in laboratories and clinical diagnosis.^2^ For example, it has been successfully applied for the detection of SARS-CoV, Middle East respiratory syndrome coronavirus (MERS-CoV), and various other viral pathogens including Zika virus (ZIKV), Influenza A virus, Dengue virus (DENV).^3-8^ After the outbreak of COVID-19, several methods and kits based on RT-PCR for the detection of SARS-CoV-2 genomic RNA have been reported.^9-10^ While RT-PCR-based methods have been widely used in COVID-19 diagnosis, their application in accurate diagnosis of viral infection and epidemic control is severely hampered by their laborious and time-consuming sample processing steps.

Quantitative extraction of nucleic acids with high purity from complex samples are the prerequisite for efficient RT-PCR assays. Low extraction efficiency might give poor signals during exponential amplification and thus result in false negative results.^11-12^ Low extraction quality, on the other hand, may contain a variety of PCR inhibitors, which gives unreliable readouts during amplification.^13-14^

In the control and diagnosis towards SARS-CoV-2 currently, silica-based spin column RNA extraction methods are widely used, in which a silica membrane or glass fiber is applied to bind nucleic acids. In these traditional methods, the samples need pre-lysis in an appropriate buffer to release nucleic acids from viral particles before binding to the column membrane and multiple centrifugation steps are required to enable binding, washing and elution of extracted nucleic acids. Additionally, as various corrosive chaotropic salts and toxic organic solvents are involved in the lysis and binding steps, several sequential washing steps are required to eliminate possible PCR inhibitory effects in the eluted products. The whole process comprises multiple centrifuging and column-transferring steps, which is laborious, time-consuming, and vulnerable to contamination or column clogging. More importantly, spin column-based approaches are not suitable for a high-throughput, automated operation. In the monitoring and control of sudden outbreaks, such as SARS-CoV-2, these traditional methods consume a large number of operators, but giving low diagnosis efficiency and high risk of cross infection. Thus, fast, convenient and automated nucleic acids extraction methods are highly desirable not just in the molecular diagnosis of SARS-CoV-2, but also in the monitoring and prevention of other infectious diseases.

As an alternative, magnetic nanoparticles (MNPs)-based extraction methods are centrifuge-free and has proven to be easy to operate and compatible to automation and high-throughput operation.^15-17^ In conventional MNPs-based methods, nucleic acids in the lysed samples can be specifically absorbed on MNPs due to various surface-modified functional groups. In the presence of magnetic fields, nucleic acids are rapidly separated from most impurities in the supernatant. After fast washing steps to eliminate trace impurities, purified nucleic acids can be further released from the surface of MNPs by elution buffer with altered ionic strength. Although much simpler and faster than spin column-based methods, most MNPs-based extraction strategies still contain multiple processing steps such as lysis, binding, washing and elution, which increases operational difficulties in real clinical diagnosis.

Herein, a further simplified and updated MNPs-based viral RNA extraction method is described for highly sensitive extraction and RT-PCR detection of viral RNA using SARS-CoV-2 pseudoviruses as models. The MNPs is synthesized via a simple one-pot approach, and functionalized with polymers carrying multi-carboxyl groups by following one-step incubation. Due to the strong interaction between carboxyl groups and nucleic acids, the poly carboxyl-functionalized MNPs (pcMNPs) enables rapid and efficient absorption of RNA molecules. In RNA extraction, one lysis/binding step and one washing step are required for nucleic acids extraction and purification from complex samples. More importantly, extracted RNA can be directed introduced into subsequent RT-PCR together with MB without elution step, which dramatically reduce operating time and risk of contamination. The fast extraction method was verified by both manual operation and automated system. Due to its simplicity, satisfactory performances and robustness, this method provides a promising alternative to decrease the labour-intensity and reduce possibility of false-negative results in current RT-PCR-based SARS-CoV-2 diagnosis.

## 3 Material and methods

### Chemicals

Iron (III) chloride hexahydrate, Iron (II) chloride tetrahydrate, ammonium hydroxide, tetraethyl orthosilicate (TEOS), (3-Aminopropyl)triethoxysilane (APTES), and dimethyl sulfoxide (DMSO) were bought from Aladdin Industrial Corporation (Shanghai, China). Ammonium hydroxide solution, isopropanol, ethylenediamine and ethanol were purchased from Sinopharm Chemical Reagent Co., Ltd. (Shanghai, China). 1,4-butanediol diacrylate, 6-amino caproic acid, sodium chloride (NaCl), sodium iodide (NaI), tri(hydroxymethyl)aminomethane (Tris), ethylene diamine tetraacetic acid (EDTA) and polyethylene glycol 8000 were obtained from Sigma-Aldrich (St. Louis, USA).

### Preparation of amino-modified magnetic nanoparticles (NH_2_-MNPs)

Bare magnetic nanoparticles (MNPs) were prepared based on a simple co-precipitation protocol as previously reported.^18^ Briefly, 3.0 g of Iron (III) chloride hexahydrate and 2.5 g of Iron (II) chloride tetrahydrate were dissolved separately in 100 mL of deionized water and degassed with nitrogen for 20 min to remove the oxygen in the solutions. Both iron solutions were then mixed in a 500 mL round-bottom flask, and 10 mL of ammonium hydroxide was added into the mixture with vigorous stirring under a nitrogen atmosphere. A rapid change of solution colour was observed from orange to black, indicating the formation of co-precipitated bare MNPs. The solution mixture was continuously stirred for another 4 h, and the resulting black products (bare MNPs) were collected with a magnet and dispersed into ethanol after washing several times with deionized water and ethanol. Subsequently, 0.3 g of as-prepared bare MNPs, 15 mL of deionized water and 12 mL of ammonium hydroxide were added into 150 mL of ethanol, followed by a continuous sonication for 30 min at room temperature. 1.2 mL of TEOS was then added dropwise into the solution mixture after sonication, and vigorously stirred for another 4 h at room temperature to allow the formation of silica layers on the surface of MNPs. Afterwards, the silica-coated MNPs were collected with a magnet and the rinsed with deionized water and ethanol for several times to remove residual TEOS. After that, 0.2 g of silica-coated MNPs were dispersed into 50 mL isopropanol, and 0.2 mL of APTES was dropwise mixed with MNPs solution. The mixture was incubated under continuous sonication for 6 h at room temperature, followed by the collection of amino-modified MNPs (NH_2_-MNPs) with a magnet and washing with deionized water and ethanol to remove free APTES. The final prepared NH_2_-MNPs were preserved in ethanol at 4 °C and the loading of -NH_2_ group on the surface was verified by Fourier transform infrared (FTIR) spectroscopy (FigureS1A).

### Synthesis and characterization of <u>p</u>oly (amino ester) with multiple <u>c</u>arboxyl groups (PC)

As shown in Scheme1 A, polymers were prepared based on the protocol reported previously with minor modification.^19^ In brief, 3.5 g of 1,4-butanediol diacrylate (17.7 mmol) and 1.5 g of 6-amino caproic acid (11.4 mmol) were firstly mixed in 10 mL of 50% (v/v) DMSO aqueous solution, followed by vigorously stirring for 12 h at 90 °C in a safe dark place. Subsequently, end-capping reaction was performed by the addition of 10% (v/v) amino solution, ethylenediamine, into the polymer/DMSO solution. The as-prepared PC polymers was final preserved at 4 °C with lightproof package. PCs were further characterized by FTIR spectroscopy (FigureS1B).

### Preparation and characterization of PC-coated NH_2_-MNPs (pcMNPs)

25 mg of NH_2_-MNPs were dispersed into 25 mL of 50% (v/v) DMSO aqueous solution and then mixed with 1.25 g of PC under sonication for 10 min. Subsequently, 2.5 mL of NaOH solution (1 M) was introduced to the mixture and sonicated for another 20 min, followed by vigorously shaking for 4 h at 37 °C and rinsing with DMSO and deionized water for several times. The obtained polymer-coated MNPs (pcMNPs) were stored in deionized water at 4 °C. The size and morphology of the pcMNPs were confirmed by transmission electron microscopy (TEM).

### Common nucleic acid extraction

The pseudovirus samples were obtained from Zeesan Biotech (Xiamen, China) and the standard samples were freshly prepared by step-wise dilution of pseudovirus in fetal calf serum purchased from Thermofisher (Massachusetts, USA) before nucleic acid extraction experiments. 200 μL of as-prepared standard samples with a known copy number of viral particles (down to 10 copies) were incubated with 400 μL lysis/binding buffer (1 M NaI, 2.5 M NaCl, 10% Triton X-100, 40% polyethylene glycol 8000, 25 mM EDTA) and 40 μg pcMNPs for 10 min at room temperature on a rotating shaker. Then, the pcMNPs-RNA complex was collected magnetically for 1 min and the supernatant was discarded. Afterwards, the complex was washed once or twice with 400 μL washing buffer (75% ethanol v/v). The purified nucleic acids were released from MNPs by incubating the complex in 50 μL of TE buffer (10 mM Tris-HCl, pH 8.0), vortexed at 55 °C bath for 5 min, and 15 μL of supernatant was transferred to subsequent RT-PCR reaction.

### Conventional Real-time reverse-transcription PCR

To quantify the amount of viral RNA captured, Real-time reverse-transcription PCR using VetMAX™-Plus One-Step RT-PCR Kit (Massachusetts, USA) and Bio-Red PCR detection system (California, USA) was applied. For conventional RT-PCR, a 30 μL of reaction solution was set up containing 15 μL of eluted supernatant, 7.5 μL of 4 × reaction buffer, 1.5 μL enzyme mixture, and optimized concentrations of primers (from 0.25 μM to 1 μM)and probe (from 0.25 μM to 1 μM). Subsequent thermal cycling was performed at 55°C for 15 min for reverse transcription, followed by 95°C for 30 s and then 45 cycles of 95°C for 10 s, 60°C for 35 s. Fluorescence readout were taken after each cycle, and the threshold cycle (Ct) was calculated by the Bio-Red analysis system based on plotting against the log10 fluorescence intensity. The oligonucleotide primers and probes were synthesized by GENEWIZ (Suzhou, China), and corresponding sequences are shown in Table 1.

**Table 1.**
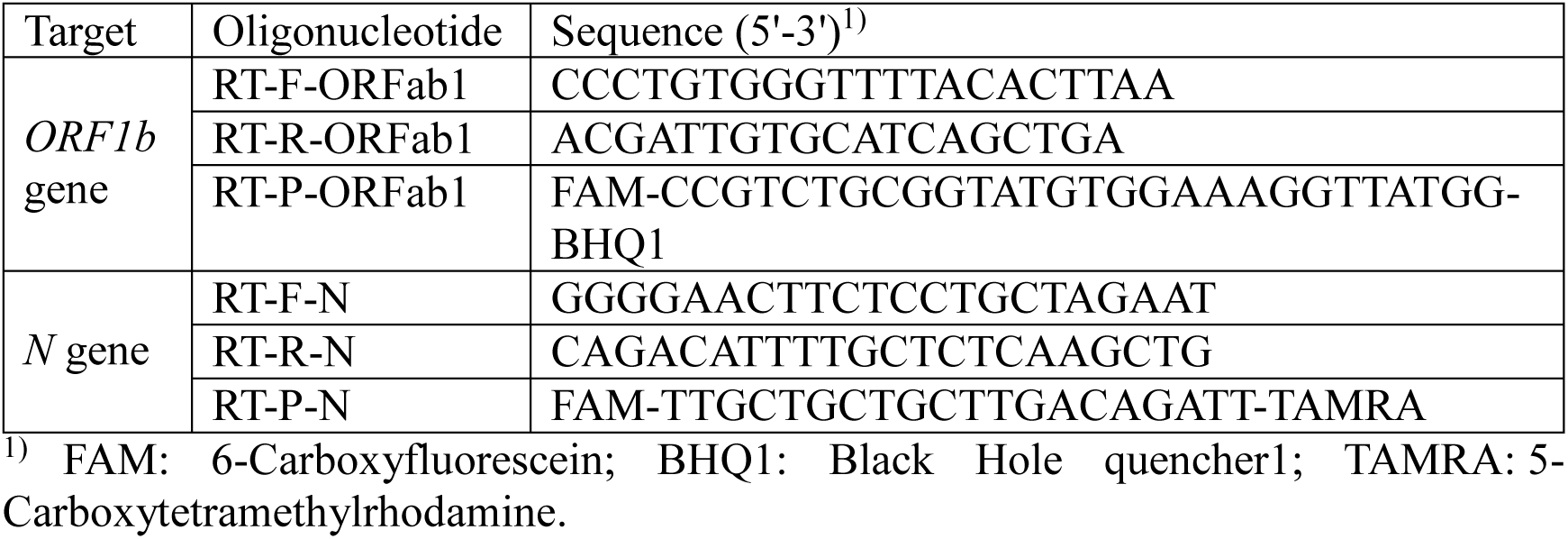
Sequences of primers and probes for RT-PCR.

### Direct RT-PCR amplification and detection of the extracted pcMNP-RNA complex

To investigate whether the extracted pcMNPs-RNA complexes can be directly used for RT-PCR, an identical extraction was carried out, in which the whole complex was transferred to the PCR tube without the elution step. Specifically, 30 μL of reaction solution composed of 15 μL of TE buffer, 7.5 μL of 4 × reaction buffer, 1.5 μL enzyme mixture, and optimal concentrations of primers (1 μM each) and probe (1 μM) was mixed with pcMNP-RNA complexes by brief vortexing, and directly transferred to the PCR tube for RT-PCR using the same procedure as abovementioned.

### Development of an automated protocol for RNA extraction

A high throughput automated RNA extraction method was adapted using a commercial NP968-C automatic nucleic acid extraction system (TIANLONG, Xi’an, China), which could simultaneously process up to 32 parallel samples in 96-well sample plates. In detail, 400 μL of lysis/binding buffer, 40 μg pcMNPs and 200 μL of the sample containing a specific number of pseudovirus particles was sequentially dispensed into Column 1 of the sample plate. Then the washing buffer and elution buffer was added separately into Column 2 or 4. Finally, the 96-well sample plate was plugged onto the matrix and RNA extraction was performed by following an optimized program (TableS1). Once the program was finished, the 96-well sample plate was removed and the 15 μL of the eluted product was analysed using the conventional RT-PCR protocol as described above.

## 4 Results and discussion

### Synthesis and characterization of pcMNPs

The pcMNPs were prepared by two steps as depicted in Scheme1B. It started with one-pot synthesis of NH_2_-MNPs using co-precipitation reaction and hydrolysis of TEOS/APTES. Once PC polymer was successfully synthesized, its reaction with NH_2_-MNPs would form pcMNPs by the Michael addition to efficiently give pcMNPs. To investigate the morphology of prepared pcMNPs, transmission electron microscopy (TEM) was used and the representative image was shown in Figure1A, suggesting a spherical morphology of the prepared pcMNPs. In addition, enlarged TEM image of pcMNPs revealed a thin silica-layer on the particle surfaces with a thickness around 3 nm, implied the successful preparation of silica-coating. By using the dynamic light scattering technique, it is observed that the prepared pcMNPs have an average diameter of 10.22 ± 2.8 nm without apparent aggregation (Figure1B). To verify the successful functionalization of synthesized PC polymers, the average zeta potentials of bare MNPs, silica-coated MNPs and pcMNPs were measured by DLS. As shown in Fig 1C, the average zeta potential changed from −4.82 mV ± 0.54 of bare MNPs to 20.57 mV ± 0.71 of silica-coated MNPs due to the NH_2_-groups introduced during silica-coating. However, functionalization of PC polymers resulted in a decrease of average zeta potential of pcMNPs to −38.67 mV ± 0.84, indicating successful polymer coating and more importantly, good dispersion of pcMNPs in the solution. Subsequently, the magnetic response property of bare MNPs, NH_2_-MNPs and pcMNPs were tested (Figure1D). The magnetically capture process was completely finished within 30 s, implying excellent paramagnetic property of the prepared pcMNPs.

**Scheme1.**
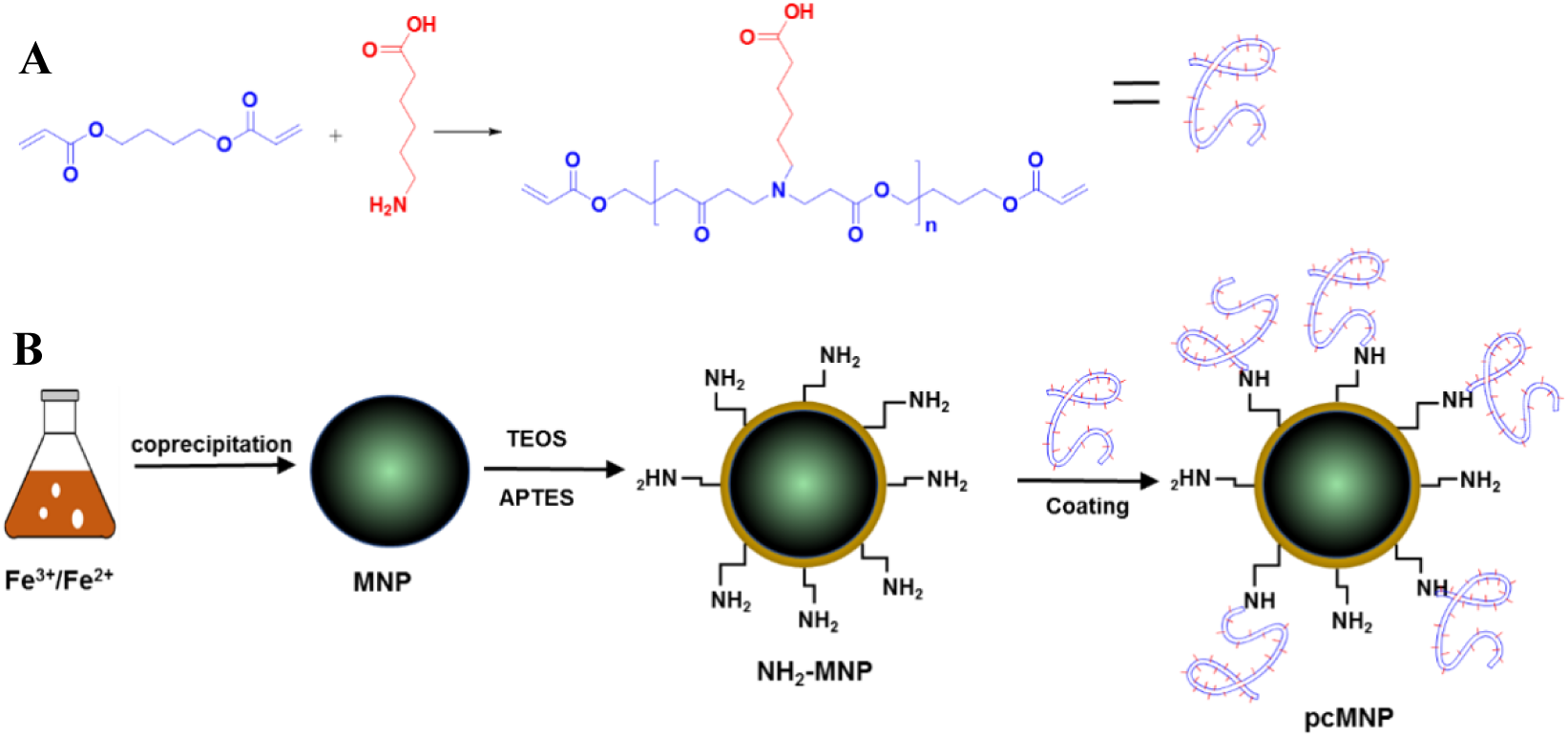
Schematic illustration for (A) the synthesis of PC polymer, and (B) the preparation of pcMNP.

**Figure1.**
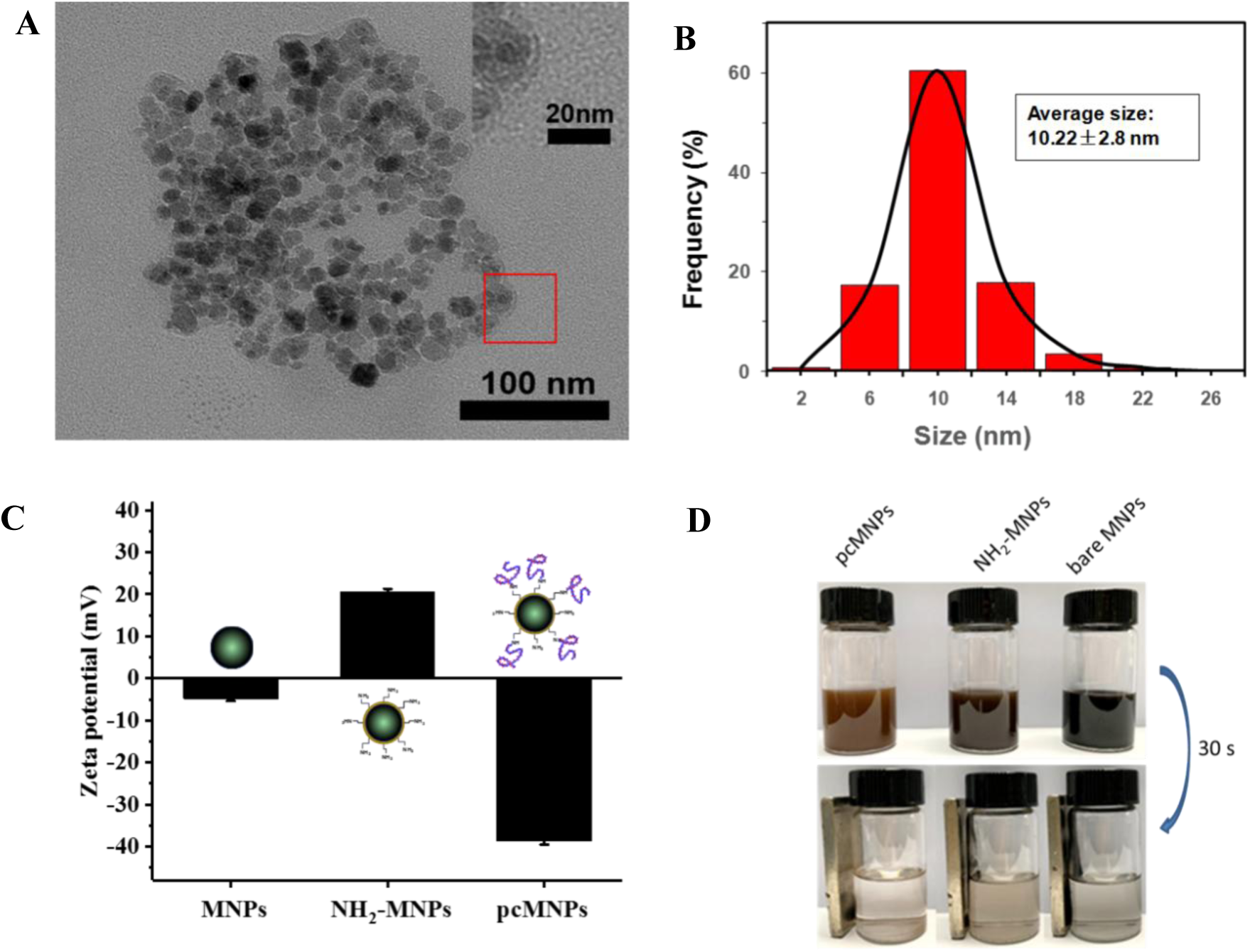
Characterization of pcMNPs. (A) TEM image and (B) size distribution of pcMNPs; The average size of nanoparticles is represented as mean ± SD of over 250 individual particles. (C) Zeta potentials of bare MNPs, NH_2_-MNPs and pcMNPs in water. (D) The magnetic response property of bare MNPs, NH_2_-MNPs and MBs in 30 s.

### RNA binding property of pxMNPs

To investigate the RNA binding property of prepared pxMNPs, RNA binding assays were performed by incubating 2 μg RNA molecules with 20 μg pcMNPs in 200 μL lysis/binding buffer. As shown in Figure2A, more than 90% of RNA was absorbed by the pcMNPs within 10 min. In contrast, significant amount RNA molecules have been observed using a commercialized RNA extraction kit under similar conditions. More importantly, no aggregation has been observed when incubating pxMNPs in a high-salt solution (50 mM NaCl), further confirmed their excellent dispersity (Figure2B).

**Figure2.**
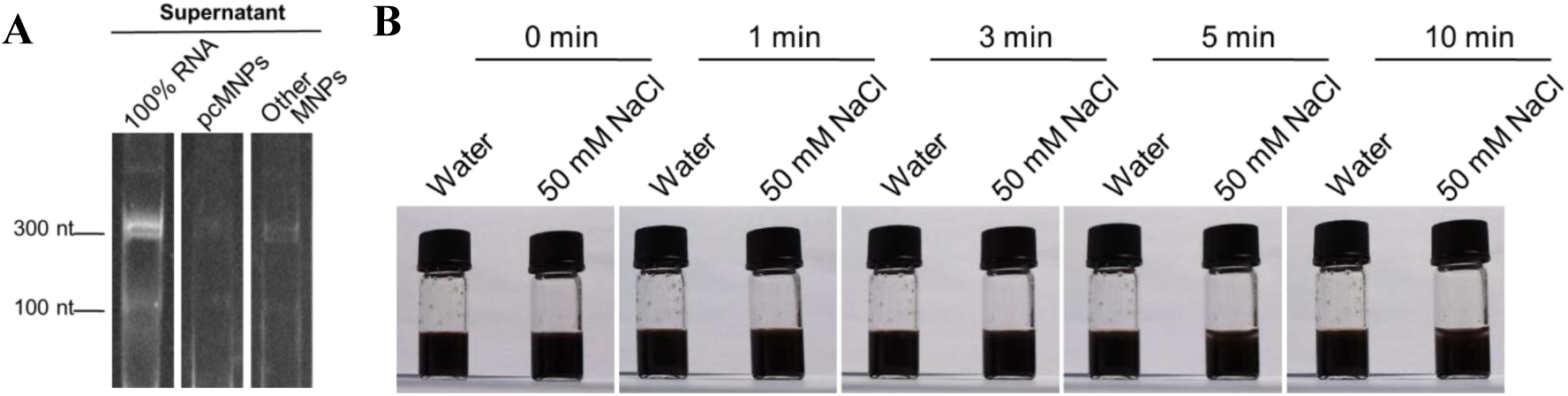
RNA binding and dispersion property of pcMNPs. (**A**) RNA binding affinity of pcMNPs analysed by native PAGE; (**B**) Dispersion stability of pcMNPs in a high-salt solution (50 mM NaCl).

### Optimization of SARS-CoV-2 viral RNA extraction and amplification

**Scheme2.**
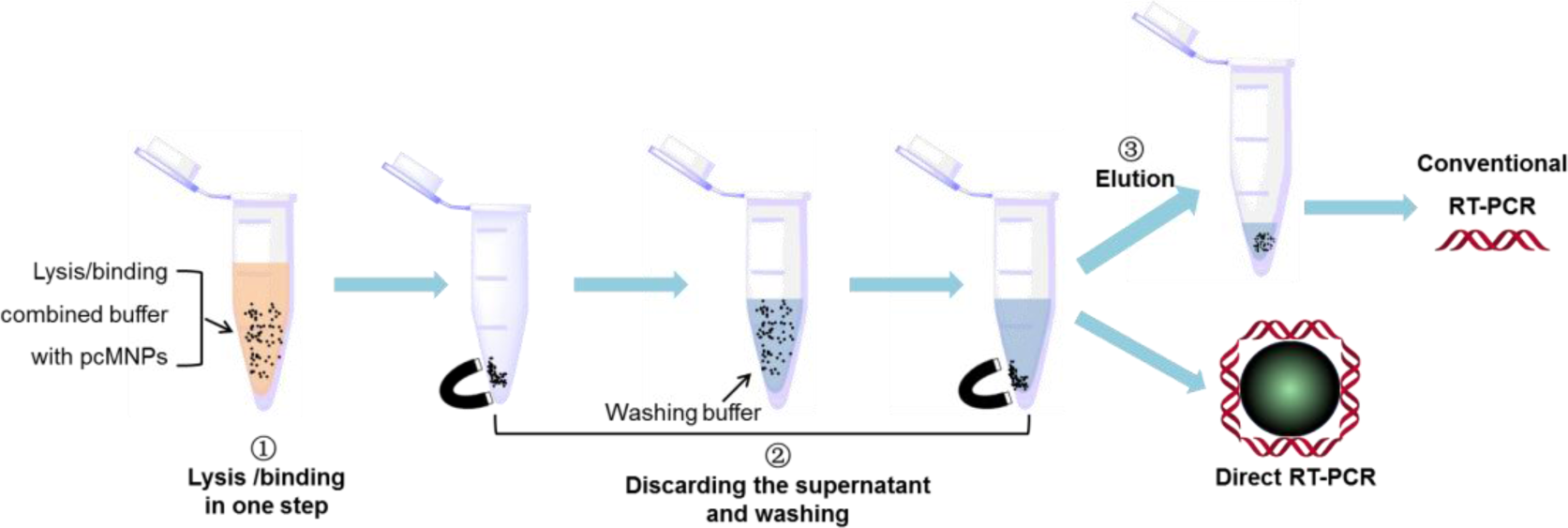
A schematic representation of the pcMNP-based viral RNA extraction method.

By following a simple lysis/binding-washing-elution protocol shown in Scheme2, 10^5^ copies of SARS-CoV-2 pseudovirus in 200 μL serum samples were extracted and subject to RT-PCR analysis. We first carried out a concentration optimization of the primer and probe to maximize the efficiency and sensitivity of SARS-CoV-2 detection. The results showed that the optimal concentrations were 1 μM and 1 μM for the primer pairs and probes respectively (FigureS2). Subsequently, the optimal amount of pcMNPs in viral RNA extraction were evaluated. Considering the high viral load in clinical samples, although 20 μg of pcMNPs were enough to extract 10^5^ copies of SARS-CoV-2 pseudovirus RNA in 200 μL serum samples (FigureS3), 40 μg pcMNPs was used in subsequent experiments.

### Applications of pcMNPs in the detection of SARS-CoV-2 viral RNA using Direct RT-PCR

Due to the excellent water dispersity of pcMNPs, we hypothesized that they can stay dispersed in the first few cycles of RT-PCR reaction, without shielding the extracted RNA molecules in aggregates from primer binding and elongation. Thus, by following the optimized conditions, performances of the pcMNPs-based RNA extraction and Direct RT-PCR was investigated. 10^5^ copies of pseudoviruses were spiked into 200 μL of serum and the extracted viral RNA were analysed by RT-PCR using eluted products (Conventional RT-PCR) or pcMNPs-RNA complexes without elution (Direct RT-PCR). In these experiments, the serum sample without pseudoviruses was used as a negative control, while the PCR reaction mixture directly spiked with 10^5^ copies of pseudoviruses was regarded as a positive control. As shown in Figure3A, in the detection of ORFlab (Open Reading Frame 1ab) region, the direct RT-PCR without elution step exhibited a slightly smaller cycle threshold (Ct) value than that measured using Conventional RT-PCR after elution. This is possibly because all extracted viral RNA was introduced to the amplification in the Direct RT-PCR, while only half of the eluted products was added for the Conventional RT-PCR. This phenomenon suggests that one advantages of the pcMNPs-based Direct RT-PCR protocol is to reduce possible false negative results, since all extracted viral RNA has been directed to the subsequent amplification without potential lost in the elution and transfer steps. Additionally, there is no detectable differences between amplification efficiencies of the positive control and Direct RT-PCR (Figure3A red and purple lines). This result suggests that our pcMNPs-based viral RNA extraction protocol not only exhibits nearly 100% RNA extraction efficiency in serum samples, but also provides high-purity products without PCR inhibitors.

**Figure3.**
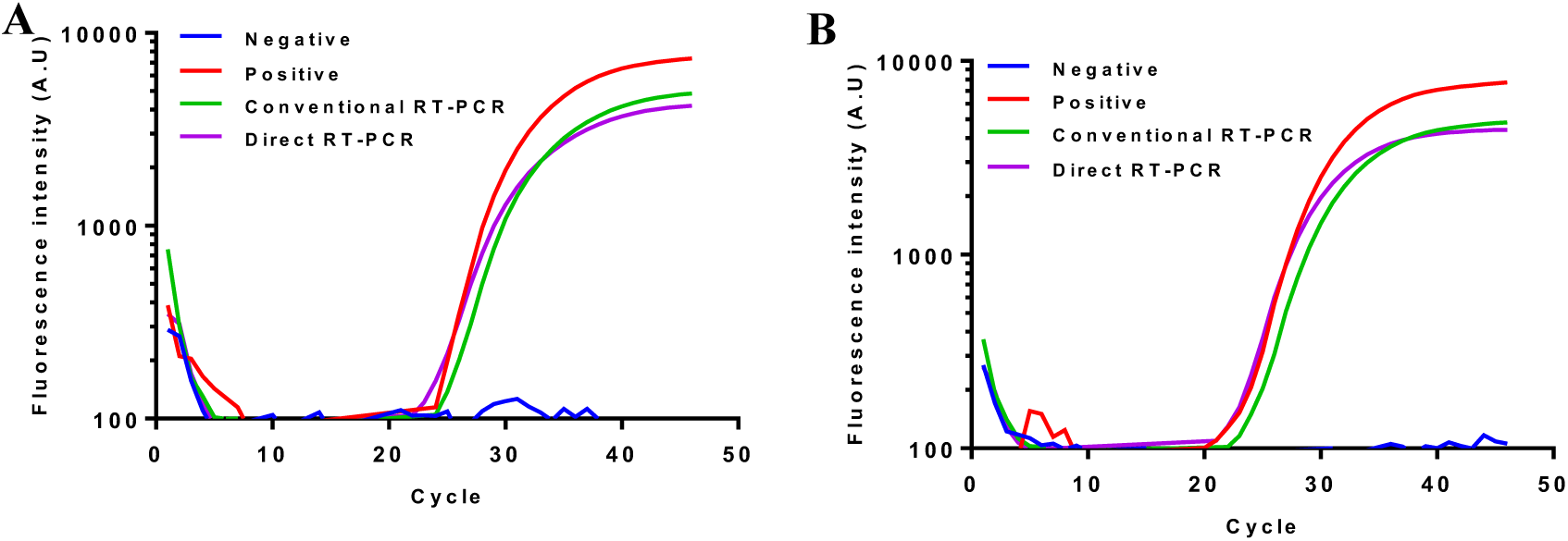
RT-PCR assays towards viral RNA extracted by different pcMNPs-based methods. RT-PCR assays amplifying (A) the ORF1ab region and (B) the *N* gene in pseudoviral RNA extracted by a manual protocol.

### Automated viral RNA extraction based on pcMNPs

As previously described, one of the most serious disadvantages of traditional column-based extraction approaches is the difficulties in automation. Because no centrifugation steps are required, MNPs-based methods allow fully automated nucleic acid purification, which is highly important in current SARS-CoV-2 diagnosis. Therefore, the feasibility of automating viral RNA extraction procedure based on our pcMNPs was subsequently evaluated. A commercialized magnetic rods-based nucleic acid purification system was used. An automated programme was set according to the manual protocol (Figure4A). During the extraction process, although the shaking pattern of magnetic rods was set at the most vigorous level, no breakage or leakage of pcMNPs were observed, since the eluted solution are colourless and transparent (FigureS4). As shown in Figure4B, the amplification curve of the automated sample is very close to that of the positive control and manually performed Direct RT-PCR samples, which suggests that our pcMNPs-based method is highly suitable for the automated high throughput viral RNA extraction.

**Figure4.**
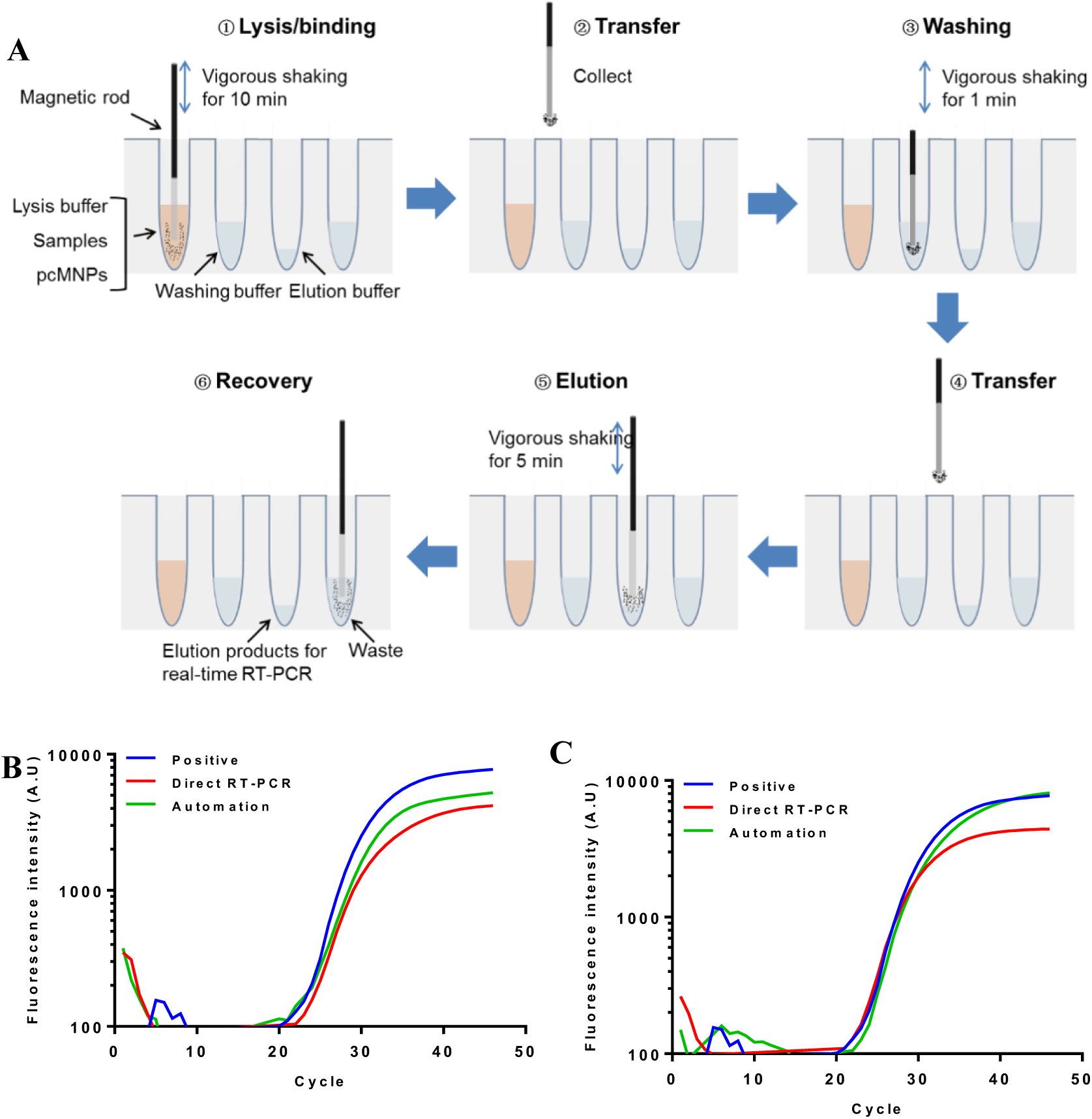
Automated viral RNA extraction based on pcMNPs. (**A**) A schematic diagram of automated extraction protocol. (B) RT-PCR assays amplifying the ORF1ab region in pseudoviral RNA extracted by automated protocol, and (B) RT-PCR assays amplifying the *N* gene in pseudoviral RNA extracted by an automated protocol.

Then, the performances of the pcMNPs-based method in extracting and detecting the *Nucleocapsid* (*N*) gene were evaluated. In agreement with the results of ORFlab region, the *N* gene assays also confirmed the high extraction efficiency and robustness of pcMNPs-based method in both manually operated Direct RT-PCR and automated protocol (Figure4C).

### Sensitivity and dynamic range of pcMNPs-based viral RNA detection

Then, the sensitivity and dynamic range of pcMNPs-based viral RNA detection method was evaluated and compared by following the Conventional and Direct RT-PCR protocols, using *N* gene carrying pseudovirus as a model. A series of standard samples containing 10^5^ to 10 copies of pseudovirus were freshly prepared by step-wise 10-fold dilution of in serum. A control serum sample (no pseudovirus) was also included as a negative control to monitor the presence of false positive signals. As shown in Figure 5A, a detection limit of 10 copies of pseudovirus was achieved using Convectional RT-PCR protocol. A strong linear relationship between the logarithm of pseudovirus particle numbers and the corresponding Ct values was found crossing over 5 orders of magnitude ranging from 10 to 10^5^ copies with a high correlation coefficient (R^2^ = 0.999) (Figure5B). Parallelly, similar detection limit and linear relationship were observed in experiments following the pcMNPs-based Direct RT-PCR protocol (Figure5C and D). Meanwhile, the differences of Ct values between neighbouring curves are approximately 3 cycles, which is very close to the theoretical 3.3 cycles for a 10-fold dilution. These results indicated that the RT-PCR amplification is highly efficient both in the presence or absence of pcMNPs. However, it is also important to note that in both cases, the negative controls sometimes gave observable amplification signals. Although their Ct values are lagged too far behind 40 cycles to be regarded as a valid positive result, this phenomenon raise concerns about possible false-positive issues, in which further optimization of primer pairs and probe might be necessary.

**Figure5.**
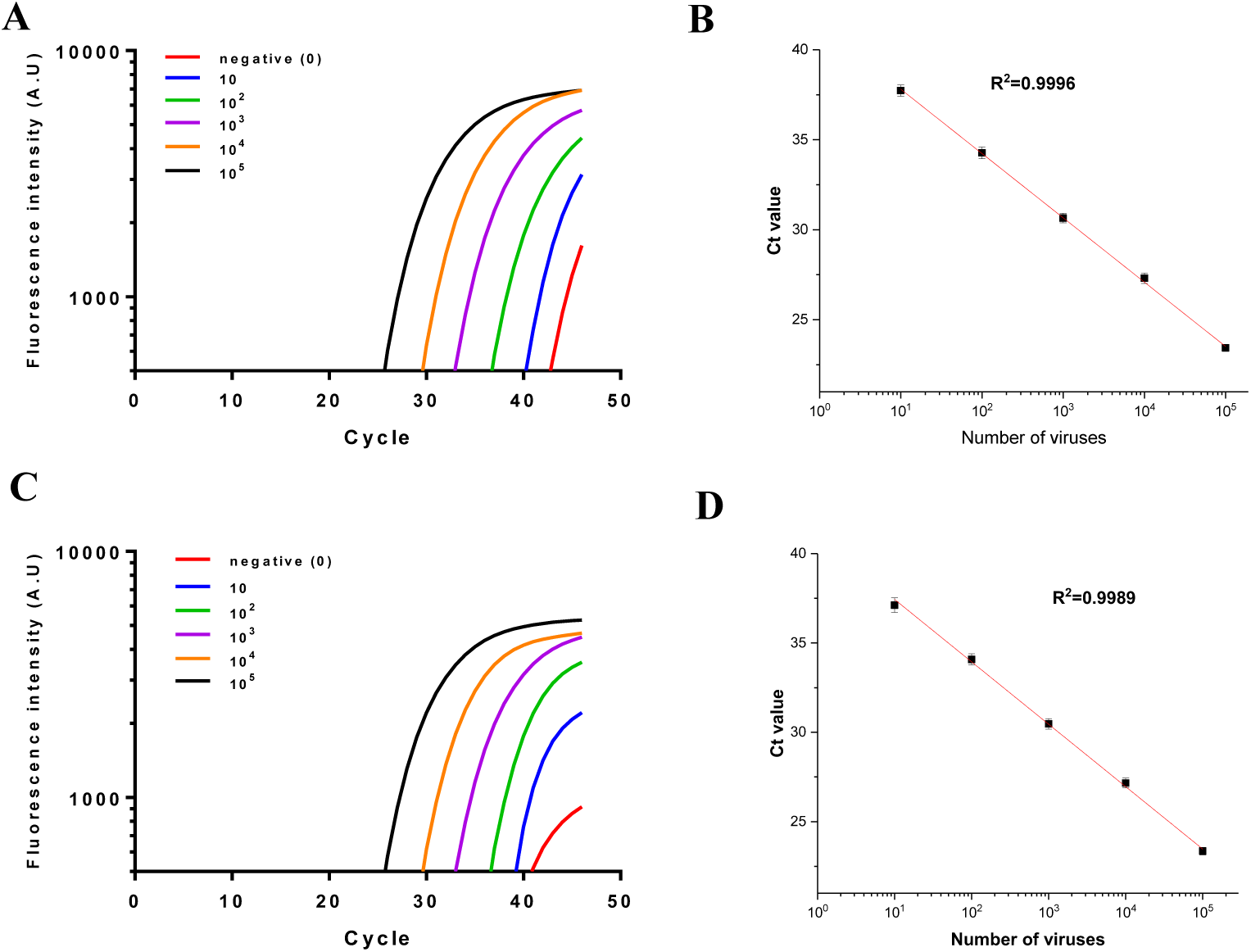
Sensitivity and dynamics range of pcMNPs-based viral RNA detection. (A) RT-PCR assays amplifying the *N* gene in pseudoviral RNA following the Conventional RT-PCR protocol, and (B) corresponding calibration curve. The red line is the linear regression fit (*R*^2^ = 0.999). (C) RT-PCR assays amplifying the *N* gene in pseudoviral RNA following the Direct RT-PCR protocol, and (D) corresponding calibration curve. The red line is the linear regression fit (*R*^2^ = 0.998).

Overall, the updated MB-based extraction method had highly extraction efficiency and compatibility of PCR amplification in any of the patterns, which dramatically simplified laborious sample processing work and was ideally suitable for RT-PCR assay of SARS-CoV-2 with a sensitivity of 10 copies at least.

## 5 Conclusions

Efficient and robust nucleic acids extraction from complex clinical samples is the first and the most important step for subsequent molecular diagnosis, but currently it is still highly labour intensive and time-consuming. For example, although the genome sequences of SARS-CoV2 have been fully revealed and various RT-PCR-based detection kits have been developed, timely diagnosis of COVID-19 is still highly challenging partially due to the lack of satisfactory viral RNA extraction strategy. In this study, a carboxyl polymer-coated MNPs, namely pcMNPs, was developed and a simple but efficient viral RNA extraction system was established for sensitive detection of SARS-CoV-2 RNA via RT-PCR. As compared with traditional column-based nucleic acids extraction methods, our pcMNPs-based method has several advantages (Table 2). Firstly, pcMNPs-based method combines the virus lysis and RNA binding steps into one, and the pcMNPs-RNA complexes can be directly introduced into subsequent RT-PCR reactions (Direct RT-PCR), which gives a dramatically simplified RNA extraction protocol. Secondly, pcMNPs have excellent viral RNA binding performances, which results in 10-copy sensitivity and the high linearity over 5 logs of gradient in SARS-CoV-2 viral RNA detection using RT-PCR. Thirdly, this method can be easily adopted in fully automated nucleic acid extraction systems without laborious optimization. Furthermore, the pcMNPs-RNA complexes obtained by this method is also compatible with various isothermal amplification methods, such as RPA and LAMP, and thus could be used in the development of POCT devices. In conclusion, due to its simplicity, robustness, and excellent performances, our pcMNPs-based method may provide a promising alternative to solve the laborious and time-consuming viral RNA extraction operations, and thus exhibits a great potential in the high throughput SARS-CoV-2 molecular diagnosis.

**Table 2.** A comparison of spin column- and pcMNPs-based extraction method in SARS-CoV-2 virus RNA extraction

| Parameter | Spin column <sup>1)</sup> | pcMNPs <sup>2)</sup> |
| --- | --- | --- |
| Complexity | Multi-step and assistant with a high-speed centrifuge | One-step and assistant with a magnet |
| Option | Manual only | Manual and automated |
| Safety | Require toxic reagents (chloroform/phenol, chaotropic salts) | No toxic reagents |
| Quality and productivity | High purity but limited productivity | High purity and high productivity |
| Elution | Require large-scale elution buffer | Directly treatment of a wide range of tested samples (food; animal; blood; pharynx; sputum and so on) |
| For RT-PCR | RNA elution products | RNA elution products or MB adsorbed RNA products without elution |
| Extraction Time for Multiple Samples | > 2 h | ~ 30 min |

<sup>1)</sup> Commercial RNA Purification: QIAGEN 52906 QIAamp Viral RNA Mini Kit; Real-time RT-PCR instrument: Roche LightCycler 480. <sup>2)</sup> This work: functional pcMNPs-based RNA extraction and real-time PCR amplification (Bio-Red).

## Reference

1. Zhu, N.; Zhang, D.; Wang, W.; Li, X.; Yang, B.; Song, J.; Zhao, X.; Huang, B.; Shi, W.; Lu, R.; Niu, P.; Zhan, F.; Ma, X.; Wang, D.; Xu, W.; Wu, G.; Gao, G. F.; Tan, W., A Novel Coronavirus from Patients with Pneumonia in China, 2019. N Engl J Med 2020, 382:727–733.

2. Espy, M. J.; Uhl, J. R.; Sloan, L. M.; Buckwalter, S. P.; Jones, M. F.; Vetter, E. A.; Yao, J. D. C.; Wengenack, N. L.; Rosenblatt, J. E.; Cockerill, F. R.; Smith, T. F., Real-Time PCR in Clinical Microbiology: Applications for Routine Laboratory Testing. Clin Microbiol Rev 2006, 19 (1), 165–256.

3. Ng, E. K. O.; Hui, D. S.; Chan, K. C. A.; Hung, E. C. W.; Chiu, R. W. K.; Lee, N.; Wu, A.; Chim, S. S. C.; Tong, Y. K.; Sung, J. J. Y.; Tam, J. S.; Lo, Y. M. D., Quantitative Analysis and Prognostic Implication of SARS Coronavirus RNA in the Plasma and Serum of Patients with Severe Acute Respiratory Syndrome. Clin. Chem. 2020, 49 (12), 1976–1980.

4. Poon, L. L. M.; Chan, K. H.; Wong, O. K.; Yam, W. C.; Yuen, K. Y.; Guan, Y.; Lo, Y. M. D.; Peiris, J. S. M., Early diagnosis of SARS Coronavirus infection by real time RT-PCR. J. Clin. Virol. 2003, 28 (3), 233–238.

5. Xu, M.-Y.; Liu, S.-Q.; Deng, C.-L.; Zhang, Q.-Y.; Zhang, B., Detection of Zika virus by SYBR green one-step real-time RT-PCR. J. Virol. Methods 2016, 236, 93–97.

6. Shisong, F.; Jianxiong, L.; Xiaowen, C.; Cunyou, Z.; Ting, W.; Xing, L.; Xin, W.; Chunli, W.; Renli, Z.; Jinquan, C.; Hong, X.; Muhua, Y., Simultaneous detection of influenza virus type B and influenza A virus subtypes H1N1, H3N2, and H5N1 using multiplex real-time RT-PCR. Appl. Microbiol. Biotechnol. 2011, 90 (4), 1463–1470.

7. Kong, Y. Y.; Thay, C. H.; Tin, T. C.; Devi, S., Rapid detection, serotyping and quantitation of dengue viruses by TaqMan real-time one-step RT-PCR. J. Virol. Methods 2006, 138 (1), 123–130.

8. Lu, X.; Whitaker, B.; Sakthivel, S. K. K.; Kamili, S.; Rose, L. E.; Lowe, L.; Mohareb, E.; Elassal, E. M.; Al-sanouri, T.; Haddadin, A.; Erdman, D. D., Real-Time Reverse Transcription-PCR Assay Panel for Middle East Respiratory Syndrome Coronavirus. J Clin Microbiol 2014, 52 (1), 67–75.

9. Corman, V. M.; Landt, O.; Kaiser, M.; Molenkamp, R.; Meijer, A.; Chu, D. K.; Bleicker, T.; Brünink, S.; Schneider, J.; Schmidt, M. L.; Mulders, D. G.; Haagmans, B. L.; van der Veer, B.; van den Brink, S.; Wijsman, L.; Goderski, G.; Romette, J.-L.; Ellis, J.; Zambon, M.; Peiris, M.; Goossens, H.; Reusken, C.; Koopmans, M. P.; Drosten, C., Detection of 2019 novel coronavirus (2019-nCoV) by real-time RT-PCR. Euro Surveill 2020, 25 (3), 2000045.

10. Chu, D. K. W.; Pan, Y.; Cheng, S. M. S.; Hui, K. P. Y.; Krishnan, P.; Liu, Y.; Ng, D. Y. M.; Wan, C. K. C.; Yang, P.; Wang, Q.; Peiris, M.; Poon, L. L. M., Molecular Diagnosis of a Novel Coronavirus (2019-nCoV) Causing an Outbreak of Pneumonia. Clin. Chem. 2020.

11. Tang, Y.; Anne Hapip, C.; Liu, B.; Fang, C. T., Highly sensitive TaqMan RT-PCR assay for detection and quantification of both lineages of West Nile virus RNA. J. Clin. Virol. 2006, 36 (3), 177–182.

12. Chan, Y. R.; Morris, A., Molecular diagnostic methods in pneumonia. Curr Opin Infect Dis 2007, 20 (2), 157–164.

13. Schrader, C.; Schielke, A.; Ellerbroek, L.; Johne, R., PCR inhibitors – occurrence, properties and removal. J Appl Microbiol 2012, 113 (5), 1014–1026.

14. Hedman, J.; Rådström, P., Overcoming Inhibition in Real-Time Diagnostic PCR. In PCR Detection of Microbial Pathogens, Wilks, M., Ed. Humana Press: Totowa, NJ, 2013; pp 17–48.

15. Pichl, L.; Heitmann, A.; Herzog, P.; Oster, J.; Smets, H.; Schottstedt, V., Magnetic bead technology in viral RNA and DNA extraction from plasma minipools. Transfusion 2005, 45 (7), 1106–1110.

16. Váradi, C.; Lew, C.; Guttman, A., Rapid Magnetic Bead Based Sample Preparation for Automated and High Throughput N-Glycan Analysis of Therapeutic Antibodies. Anal. Chem. 2014, 86 (12), 5682–5687.

17. Riemann, K.; Adamzik, M.; Frauenrath, S.; Egensperger, R.; Schmid, K. W.; Brockmeyer, N. H.; Siffert, W., Comparison of manual and automated nucleic acid extraction from whole-blood samples. J Clin Lab Anal 2007, 21 (4), 244–248.

18. Song, W.; Su, X.; Gregory, D. A.; Li, W.; Cai, Z.; Zhao, X., Magnetic Alginate/Chitosan Nanoparticles for Targeted Delivery of Curcumin into Human Breast Cancer Cells. Nanomaterials 2018, 8 (11), 907.

19. Sunshine, J.; Green, J. J.; Mahon, K. P.; Yang, F.; Eltoukhy, A. A.; Nguyen, D. N.; Langer, R.; Anderson, D. G., Small-Molecule End-Groups of Linear Polymer Determine Cell-type Gene-Delivery Efficacy. Adv Mater 2009, 21 (48), 4947–4951.

