## Supporting information for "A simple magnetic nanoparticles-based viral RNA extraction method for efficient detection of SARS-CoV-2"

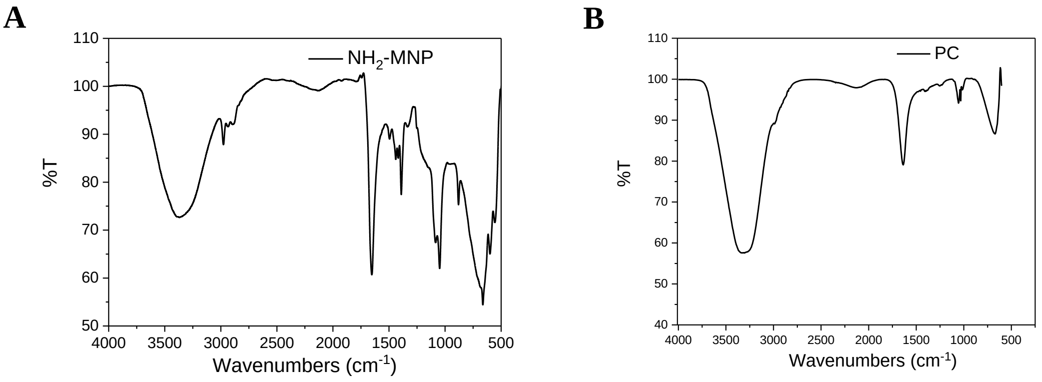


**FigureS1 Characterization of $NH_2$-MNP (A) and PC (B) by FTIR spectroscopy.**

| Step | Column | Temperature | Reaction time | Frequency | Magnetically capture time | Magnetic frequency |
|---|---|---|---|---|---|---|
| Lysis/ Binding | 1 | 96℃ | 10 min | Fast | 1min | Slow |
| Wash1 | 2 | - | 1 min | Medium | 1min | Slow |
| Wash2 | 3 | - | 1 min | Medium | 1min | Slow |
| Elution | 6 | 86℃ | 5 min | Fast | 1min | Slow |
| Recovery | 2 | - | 15 s | Fast | - | - |

**TableS1 The program of NP968-C automatic nucleic acid extraction system.**

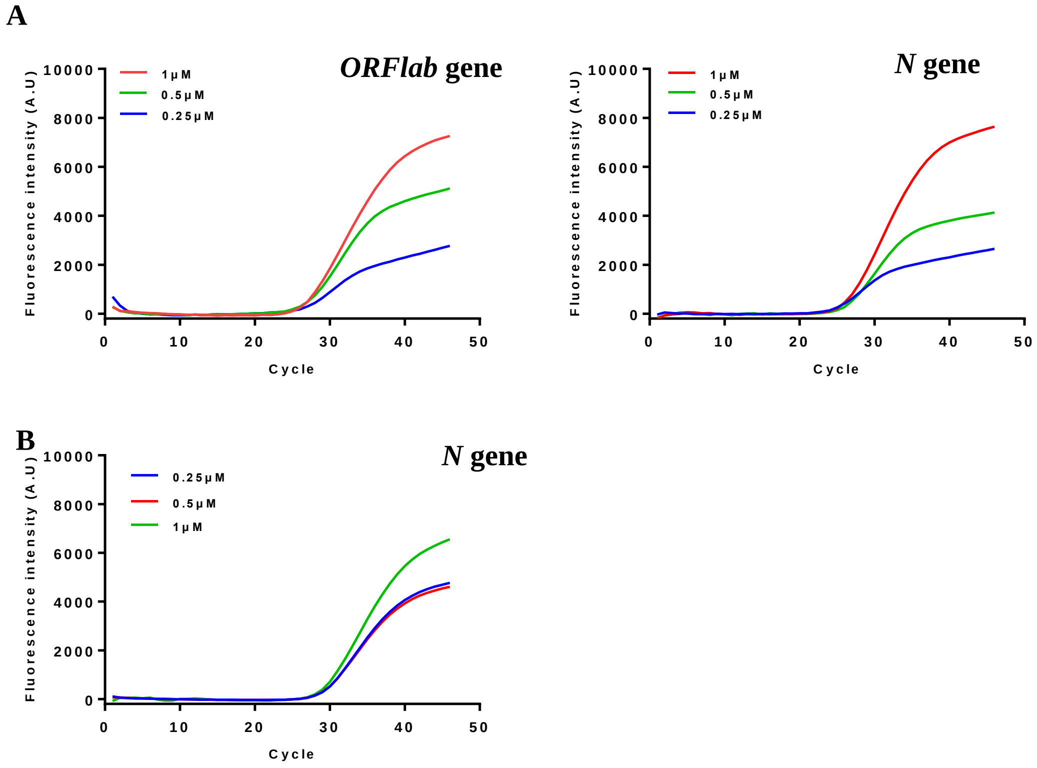


**FigureS2 Optimization of the concentrations of primer pairs and TaqMan probes.** (A) RT-PCR assays amplifying ORF1ab region and *N* gene region in the pseudoviral RNA with different primer pair concentrations. (B) RT-PCR assays amplifying *N* gene region in the pseudoviral RNA with different TaqMan probe concentrations.

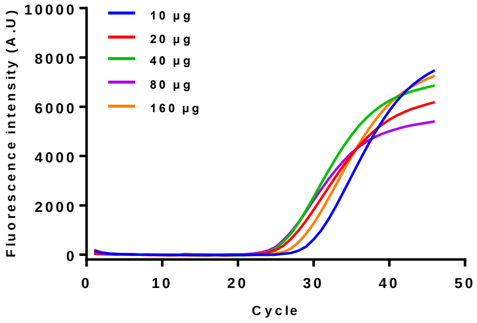


**FigureS3 Optimization towards the amount of pcMNPs used for extraction.**

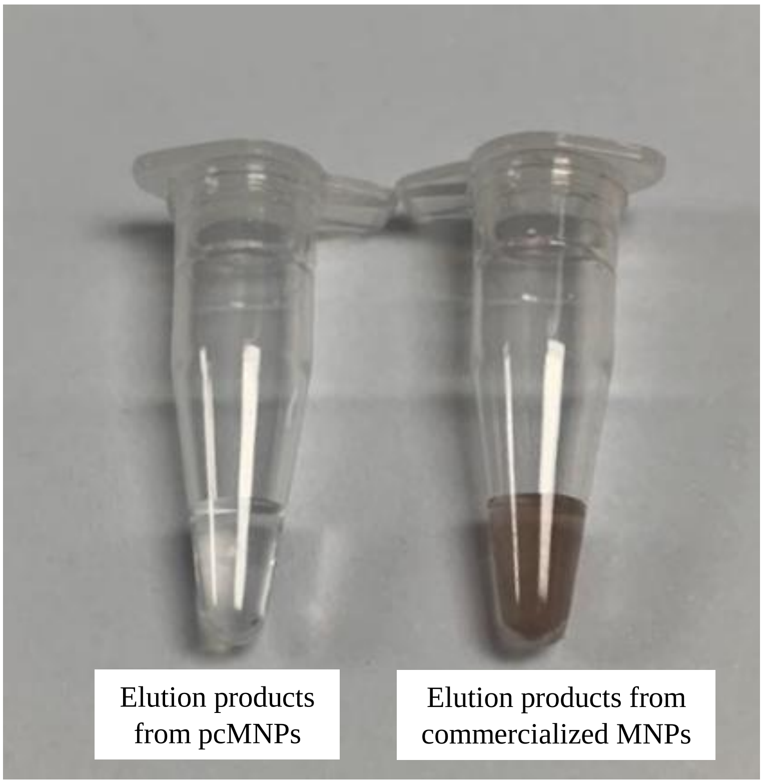


**FigureS4 Breakage or leakage of commercially available MNPs under automation.**
